# Architecture of parallel adaptation to freshwater in multiple populations of threespine stickleback

**DOI:** 10.1101/381723

**Authors:** Nadezhda V. Terekhanova, Anna E. Barmintseva, Alexey S. Kondrashov, Georgii A. Bazykin, Nikolai S. Mugue

**Affiliations:** Skolkovo Institute of Science and Technology, Skolkovo, Russia; Sector for Molecular Evolution, Institute for Information Transmission Problems of the RAS (Kharkevich Institute), Moscow, Russia; Laboratory of Molecular genetics, Russian Federal Research Institute of Fisheries and Oceanography, Moscow, Russia; Department of Ecology and Evolutionary Biology and Life Sciences Institute, University of Michigan, Ann Arbor, Michigan, United States of America; M. V. Lomonosov Moscow State University, Moscow, Russia; N. K. Koltsov Institute of Developmental Biology RAS, Moscow, Russia

## Abstract

Threespine sticklebacks adapted to freshwater environments all over the Northern Hemisphere. This adaptation involved parallel recruitment of freshwater alleles in clusters of closely linked sites, or divergence islands (DIs). However, it is unclear to what extent the DIs involved in adaptation and the alleles within them coincide between populations adapting to similar environments. Here, we examine 10 freshwater populations of similar ages from the White Sea basin, and study the repeatability of patterns of adaptation in them. Overall, the 65 detected DIs tend to reside in regions of low recombination, underlining the role of reduced recombination in their establishment. Moreover, the DIs are clustered in the genome to the extent that is not explainable by the recombination rate alone, consistent with the divergence hitchhiking model. 21 out of the 65 DIs are universal; i.e., the frequency of freshwater alleles in them is increased in all analyzed populations. Universal DIs tend to have longer core region shared between populations, and the divergence between the marine and the freshwater haplotypes in them is higher, implying that they are older, also consistently with divergence hitchhiking. Within most DIs, the same set of sites distinguished the marine and the freshwater haplotypes in all populations; however, in some of the DIs, the genetic architecture of the freshwater haplotype differed between populations, suggesting that they could have been established by soft selective sweeps.

## Introduction

The wide range of threespine stickleback *Gasterosteus aculeatus* encompasses both coastal seas and freshwater bodies in the Northern Hemisphere. Its marine populations can rapidly evolve adaptations to freshwater lakes and streams (Jones et al. 2012; Marques et al. 2016). A newly formed freshwater lake may soon become colonized by individuals of the marine ancestry that give rise to the freshwater population. This process often takes place independently in different lakes (Colosimo et al. 2005). If the connection between the lake and the sea persists, there gene flow between the ancestral marine and the derived freshwater population may remain (Roesti et al. 2015; Pedersen et al. 2017).

Marine and freshwater environments of threespine stickleback are drastically different; in particular, different salinity, parasites, and predators exert divergent selective pressures on its population. A salient phenotypic difference between the marine and the freshwater individuals is that only the former possess a complete row of bony plates along the lateral line. Genes that are likely to be involved in the differential adaptation to the two environments include osmoregulatory genes ATPase 1 alpha-subunit *ATP1A1* and V-type ATPase *VMA21* (DeFaveri et al. 2011; Jones et al. 2012), genes that encode important components of the immune system such as *GARP* (Colosimo et al. 2005; Robertson et al. 2017), and genes responsible for the embryogenesis *WNT7B* (Jones et al. 2012) and development of the lateral plates and gill rakers, *EDA* and *EDAR* (Colosimo et al. 2005; Jones et al. 2012; Glazer et al. 2014). A number of genes, such as *MUC-like* genes (Jones et al. 2012; Seear et al. 2015), may play a role in the pre-zygotic isolation between freshwater and marine individuals.

One could think that the adaptation of threespine stickleback populations to the radically new freshwater environment occurs independently in every case, and relies on *de novo* mutations. However, this adaptation often proceeds very fast, over the course of several decades (Terekhanova et al. 2014; Lescak et al. 2015; Marques et al. 2018), with some changes becoming detectable even sooner (Barrett et al. 2008). Clearly, such rapid evolution cannot depend on *de novo* mutations (Orr 2017) and must rely primarily on the standing genetic variation (Schluter and Conte 2009; Karasov et al. 2010; Matuszewski et al. 2015). Indeed, marine populations of threespine stickleback harbor, at low frequencies, alleles that confer adaptation to freshwater (Schluter and Conte 2009), presumably due to gene flow from the coastal freshwater populations (Bassham et al. 2018). Although such alleles must be deleterious under the unsuitable environment, the resulting selection is not strong enough to eliminate them immediately (Bassham et al. 2018). As a result, the sets of genetic markers that distinguish derived freshwater populations from the ancestral marine population (marker SNPs) are strongly similar across different lakes (Hohenlohe et al. 2010; Jones et al. 2012; Terekhanova et al. 2014).

Overall, marine and freshwater genotypes are very similar to each other; however, there is a number of genomic regions where their divergence is quite high. If only such regions, known as islands of divergence (DIs) (Turner et al. 2005; Feder and Nosil 2010), are taken into account, all threespine stickleback populations become subdivided into marine and freshwater clades (Colosimo et al. 2005; Nelson and Cresko 2018). DIs can be identified by their enrichment with the marker SNPs that have substantially different allele frequencies in the marine and freshwater populations. DIs have been observed in a number of species that recently evolved adaptations to new environments (Ellegren et al. 2012; Jones et al. 2012; Renaut et al. 2013).

DIs are often characterized by reduced recombination (Feulner et al. 2015; Samuk et al. 2017). They are also characterized by increased linkage disequilibrium (Hohenlohe et al. 2011; Larson et al. 2016), which could be partially due to recent selective sweeps or background selection (Ellegren et al. 2012; Feulner et al. 2015). Local selection sweeps decrease the interpopulation gene flow in the genomic region around the target of positive selection (Smith and Haigh 1974; Barton 2000). After the allele replacement is over, the length of the affected region decreases with time due to recombination, as long as some migration between the populations persists. Of course, if a gene involved in local adaptation is situated within an inversion, the whole inversion may become a DI (Kirkpatrick 2010; Sodeland et al. 2016). Still, some DIs can appear in the regions of usual recombination if divergent selection is strong (Feder et al. 2012). A DI can emerge around a single locus that undergoes local positive selection; however, multiple, tightly linked targets of positive selection within a DI are also possible, and it may be difficult to distinguish these two possibilities.

The distribution of marker SNPs within a DI is non-uniform, and, as long as some recombination takes place, their density is higher close to the target(s) of divergent selection. Moreover, the density of maker SNPs increases with time since the incipience of such selection. DIs can be ancient (Ma et al. 2018; Nelson and Cresko 2018), and the DIs found in threespine stickleback originated, on average, ~6 Mya and were shaped by the recurrent action of divergent selection after that (Nelson and Cresko 2018). In the course of their long history, these DIs accumulated many marker SNPs that distinguish the marine and the freshwater haplotypes and, in some cases, inversions which suppress recombination between them. When an inversion is present, its boundaries may coincide with the boundaries of the corresponding DI, in which case the density of marker SNPs may be uniform across the whole DI (Jones et al. 2012; McGaugh and Noor 2012; Nadeau et al. 2012; Sodeland et al. 2016).

A DI can evolve through divergence hitchhiking, which leads to fixations of new slightly beneficial mutations in only one of the alternative haplotypes, in the vicinity of loci where two different alleles have already been selected (Via 2009; Feder et al. 2012). Theory predicts that divergence hitchhiking leads to extension of DIs (Feder and Nosil 2010; Feder et al. 2012), which is aided by the presence of structural variants that suppress recombination (Flaxman et al. 2013; Yeaman 2013; Yeaman et al. 2016). Data analyses provide conflicting estimates of the impact of divergence hitchhiking on the evolution of DIs (Hohenlohe et al. 2011; Renaut et al. 2012; Via 2012; Burri et al. 2015; Feulner et al. 2015).

Data on genotypes of multiple freshwater populations (Hohenlohe et al. 2010; Jones et al. 2012; Terekhanova et al. 2014) show that a large proportion of marker SNPs are present in many, or even all of them. Thus, evolution of adaptations to freshwater proceeds through assembling of “pre-cast bricks” of freshwater-adapted haplotypes which are a part of the standing genetic variation in the marine populations (Terekhanova et al. 2014). This model predicts that these adaptations may evolve through soft selective sweeps (Orr and Betancourt 2001; Messer and Petrov 2013; Garud et al. 2015). Some data support this prediction (Roesti et al. 2014; Bassham et al. 2018), although the prevalence of soft selective sweeps has not been examined systematically.

Here, we study DIs that exist in 14 freshwater populations of threespine stickleback that independently evolved from the marine population of the White Sea. Examination of the genomic sequences of DIs in multiple populations that adapted in parallel allows us to explore the interplay between the processes which are involved in their formation and to make inferences about contributions of individual loci into this adaptation.

## Results

### Genomic distribution of DIs

We performed pooled sequencing of between 8 and 24 individuals from each of the ten relatively old freshwater populations and of the four populations of recent origin (table 1 and fig. 1A), six of which had been analyzed previously (Terekhanova et al. 2014). We identified a total of 180,249 marker diallelic SNPs where an allele with a frequency below 0.2 in the ancestral marine population reaches a frequency above 0.8 in at least one of the ten older freshwater populations. These marker SNPs are clustered into 65 DIs (fig. 2 and table S1, see Materials and Methods). The overall divergence between marine and freshwater populations within these DIs is almost 2.5 times higher than in the rest of the genome (mean Dxy over 10 comparisons: 0.0062 within DIs and 0.0026 outside DIs; fig. 1B).

**Table 1.** Description of the locations of populations studied. The sequencing reads from the three marine populations were pooled together. The populations marked with an asterisk have been analyzed previously (Terekhanova et al. 2014).

| Sample | ID | Description | Geographic location | Number of individuals | Number of reads | # of reads properly paired | Mean coverage |
| --- | --- | --- | --- | --- | --- | --- | --- |
| White Sea, WSBS | MAR | Marine | 66°57.04'N, 33°10.40'E | 12 | 415,169,461 | 365,058,887 | 90 |
| Nilma* |  | Marine | 66°30.45'N, 33°7.68'E | 16 |  |  |  |
| Ershovskoye*<br>(anadromous) |  | Marine | 66°32.21'N, 33°3.62'E | 10 |  |  |  |
| Lobaneshskoye* | LOB | Freshwater, older | 66°33.64'N, 33°13.45'E | 8 | 120,452,652 | 81,595,168 | 20 |
| Mashinnoye* | MAS | Freshwater, older | 66°17.74'N, 33°21.82'E | 10 | 92,891,151 | 64,737,645 | 16 |
| Canon | CAN | Freshwater, older | 66°16.69'N, 34°13.33'E | 24 | 202,051,636 | 190,057,977 | 45 |
| Lake Nilma | LN | Freshwater, older | 66°49.15'N, 33.09.75'E | 16 | 185,364,807 | 171,676,383 | 41 |
| Son | SON | Freshwater, older | 66°17.64'N, 34°14.73'E | 24 | 76,682,927 | 73,382,541 | 25 |
| Mashinnoye-3 | MAS3 | Freshwater, older | 66°29.85'N, 33°34.41'E | 12 | 44,514,740 | 41,667,387 | 15 |
| Kumyazh'i | KUM | Freshwater, older | 66°56.24'N, 33°32.63'E | 24 | 156,410,799 | 147,101,391 | 35 |
| Ogorognoye | OG | Freshwater, older | 66°56.80'N, 33°21.07'E | 20 | 89,328,329 | 79,583,953 | 27 |
| Belaya Guba | BG | Freshwater, older | 66°91.08'N, 32°45.83'E | 20 | 69,020,533 | 64,178,878 | 22 |
| Voron'ye | VOR | Freshwater, older | 66°95.04'N, 32°41.90'E | 20 | 73,677,728 | 63,567,981 | 21 |
| Malysh* | MAL | Freshwater, 33 yrs | 66°18.27'N, 33°25.27'E | 20 | 308,787,163 | 270,452,403 | 68 |
| Goluboy* | GOL | Freshwater, 33 yrs | 66°17.20'N, 33°23.29'E | 20 | 247,299,313 | 165,873,385 | 41 |
| Ershovskoye*<br>(residential) | ER | Freshwater, ~30 yrs | 66°32.21'N, 33°3.62'E | 12 | 255,561,219 | 204,988,298 | 51 |
| Martsy* | MART | Freshwater, ~250 yrs | 66°35.95'N, 33°15.35'E | 10 | 126,624,880 | 77,327,734 | 19 |

**Fig. 1.**
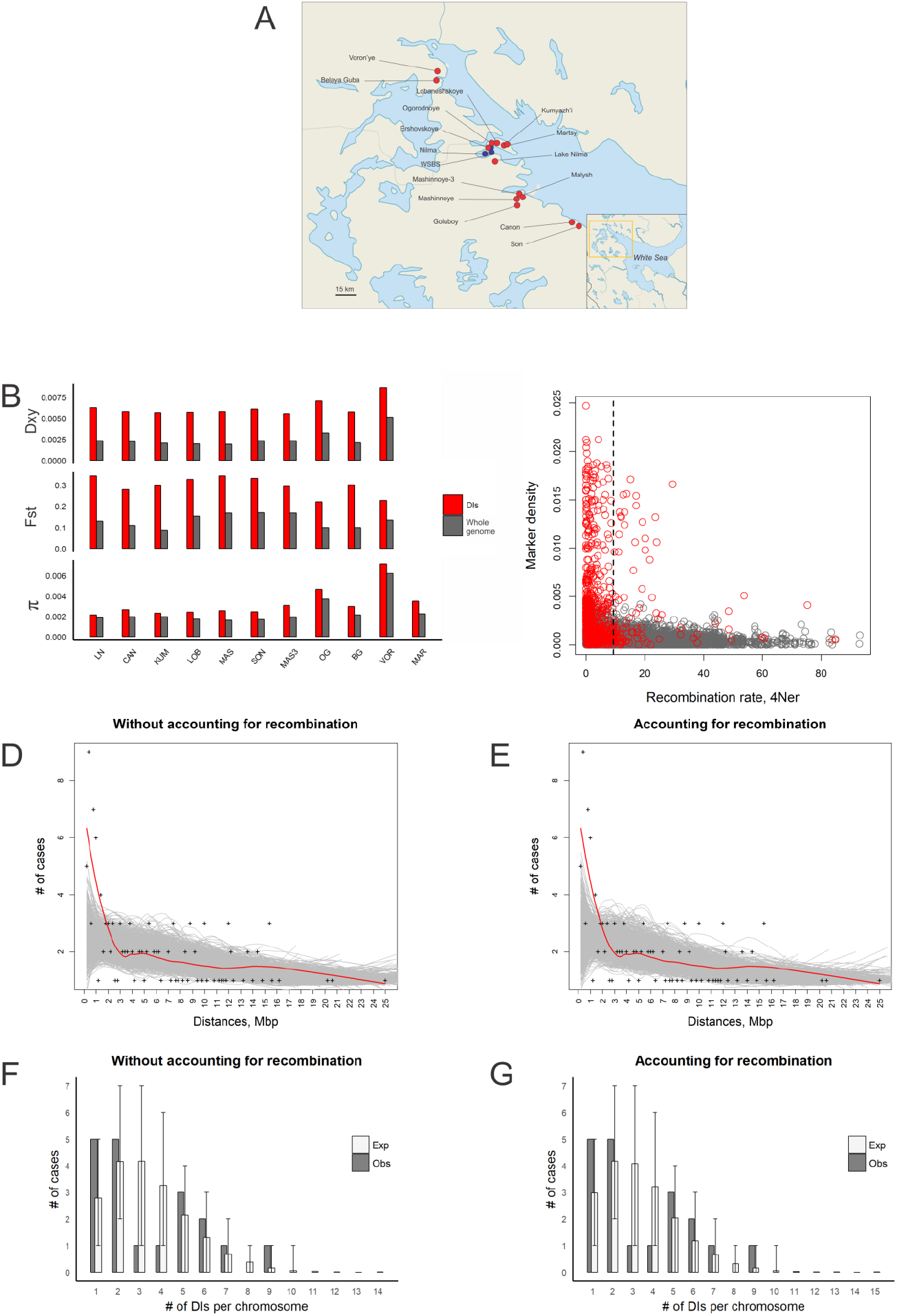
Divergence in the freshwater populations. **(A)** Map showing the locations of populations studied. **(B)** Dxy, Fst and π calculated between one marine and ten freshwater populations studied. **(C)** Marker SNP density plotted against recombination rate in 10 Kb genomic windows. Red color denotes windows located inside identified DI regions; dark grey, all other genomic segments. Black dashed line, the average recombination rate across the genome. **(D-E)** Distribution of distances between all pairs of DIs located on the same chromosome. Each cross denotes the number of pairs of DIs falling into a particular 200 Kb distance bin (horizontal axis) from each other. Lines, loess smoothing (span = 0.5) for actual data (red) or for each of the 1000 reshuffling trials (grey) without **(D)** and with **(E)** accounting for local recombination rate. **(F-G)** Distribution of the numbers of DIs per chromosome without **(F)** and with **(G)** accounting for the local recombination rate. Expected numbers are calculated from the 1000 permutation trials.

**Fig. 2.**
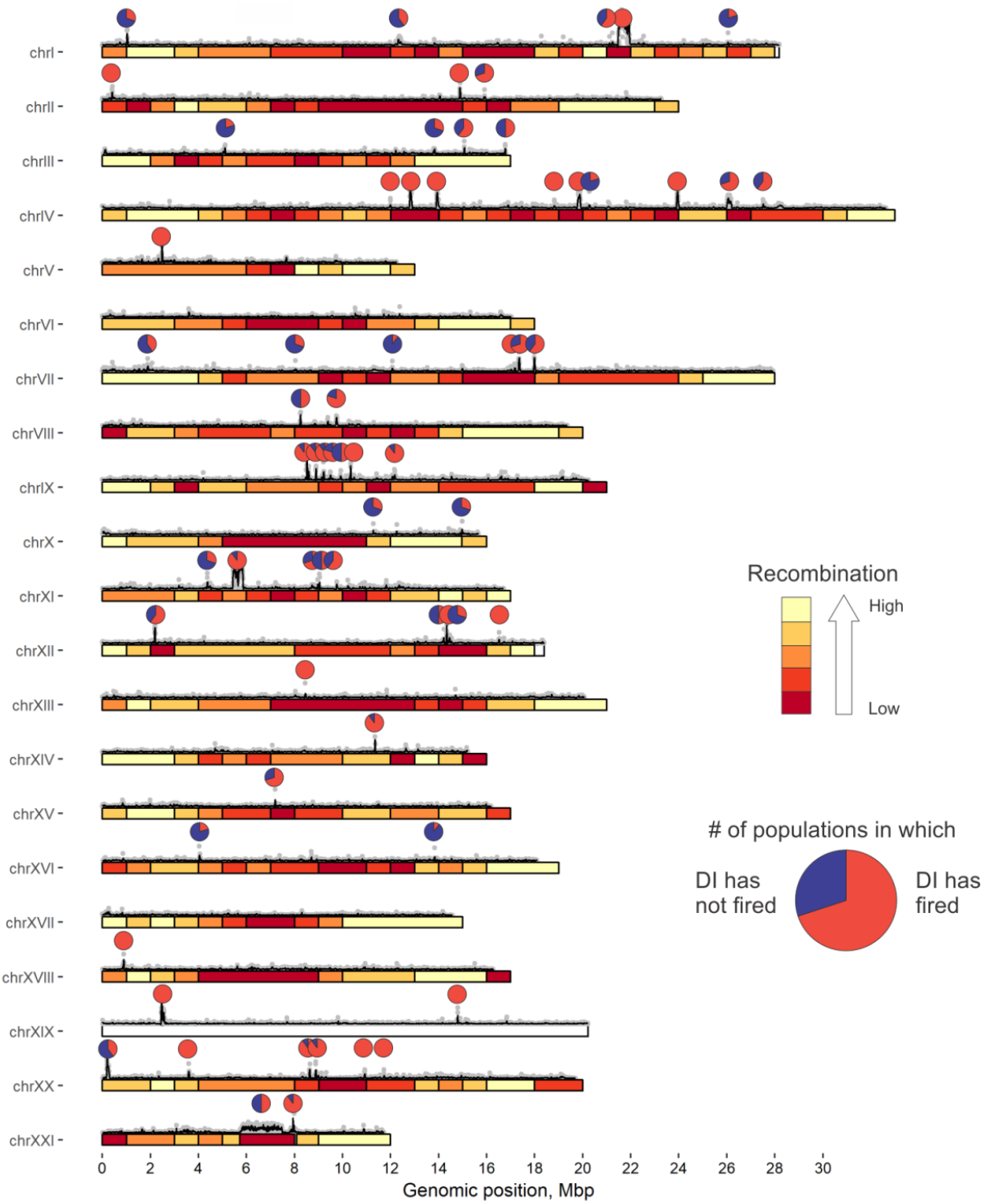
Genomic positions of DIs. The colors from red to yellow correspond to the 5 bins of recombination rate from low to high (average values for 1 Mb genomic windows). For each identified DI, the pie chart shows DI pervasiveness, i.e., the fraction of the populations (out of 10) in which this DI has “fired”, so that the marker SNPs carry the freshwater alleles at mean frequency of > 0.5. Above the bars: grey dots, the numbers of markers in each 5 Kb window; black lines, their smoothing by loess function with the span = 0.0005.

The DIs are characterized by low rates of recombination (mean coalescent-based population recombination rates 4*N_e_r* (Feulner et al. 2015) for DIs and for the whole genome are 6.63 and 8.24, respectively; one-sided Wilcoxon test P = 1.23×10^−4^). The boundaries of three DIs I-4, XI-2, XXI-1 match the boundaries of known inversions, in line with previous findings (Jones et al. 2012). However, even within the non-inversion DIs, the recombination rate (6.73) is lower than the genome average (one-sided Wilcoxon test P = 2.0× 10^−4^). Still, some DIs recombine fast: the recombination rates within 11 DIs are above the genome average by factors of up to 7.7 (fig. 1C and table S2).

The DIs are clustered within chromosomes. In 30 DI pairs, the two DIs are located within 1 Mb of one another, although only 10.6 such pairs are expected if the DIs were distributed over the genome randomly (P < 0.001, fig. 1D). This difference remains significant even when the randomization procedure takes into account the heterogeneity of the recombination rate between genomic regions (30 vs. 11.3, P < 0.001, fig. 1E). Thus, within-chromosome clustering of DIs is not explainable by variation of recombination rates. The variance of the numbers of DIs per individual chromosome was also higher than expected if they were distributed randomly, although the difference was of minor significance (2.411 observed vs. 1.879 expected, P = 0.041, fig. 1F). We see the same pattern when the recombination rate is controlled for (2.411 observed vs. 1.875 expected, P = 0.053, fig. 1G).

### Differences in architecture of adaptation between freshwater populations

While an increase in the frequency of freshwater alleles within a DI in a freshwater population denotes adaptation to freshwater, we asked to what extent this adaptation is repeatable between freshwater populations. Let us say that a DI has “fired” in a particular freshwater population if the mean frequency of freshwater alleles at marker SNPs across at least the 20% of this DI is above 0.5 (figs. 3A–B). Out of the 65 DIs, 45 (69%) fired in an average population. 21 (32%) DIs fired in each population (hereafter, universal DIs), and 49 (75%) fired in at least half of the populations. Only 2 (3%) DIs fired in just one population, and both of them fired in the SON population (table S1). We define DI pervasiveness as the proportion of populations in which it has fired (fig. 2).

DIs residing within known inversions were pervasive and fired on average in eight populations (table S2). Among non-inversion DIs, universal DIs differed from the remaining DIs in several respects. They had higher mean frequencies of the freshwater allele across population where they fired (0.86 vs. 0.79, one-sided Wilcoxon rank sum test P = 5.6×10^−3^), longer core regions shared between populations (see below; 40.1 Kb vs. 29.4 Kb, one-sided Wilcoxon rank sum test P = 0.046), and higher density of marker SNPs in the core region (0.0058 vs. 0.0039, one-sided Wilcoxon rank sum test P = 0.024).

In order to investigate whether some pairs of DIs tend to fire in the same populations, we measured, for every pair of DIs, the correlation between their mean freshwater allele frequencies across the 10 older freshwater populations. We found a total of 7 significant correlations, 6 of which were positive and the remaining one negative (table S3). DIs from significantly correlated pairs harbor several genes involved in immune response and reproduction (tables S3 and S4).

### Genetic architecture and evolutionary history of individual DIs

Availability of multiple freshwater populations allowed us to study the reproducibility of the allelic composition of a DI between populations. In line with the “precast bricks” model (Terekhanova et al. 2014), one can assume that at a particular DI, the same freshwater-adapted haplotype was recruited and spreads in every freshwater population in which this DI has fired. In this simplest case, a DI is the result of just one ancestral “hard” sweep (Pennings and Hermisson 2006) which gave rise to the freshwater-adapted haplotype. Alternatively, multiple freshwater-adapted haplotypes could arise at a given DI in the metapopulation of sticklebacks (Bassham et al. 2018) by the process of “soft” sweep (Pennings and Hermisson 2006; Barrett and Schluter 2008; Messer and Petrov 2013; Hermisson et al. 2017). If these haplotypes have survived to the present, we may observe that the same DI would have different marker composition in different freshwater populations.

First, we characterized the diversity of haplotypes within each DI in terms of their allelic composition. In most DIs the sets of marker SNPs that distinguish different freshwater populations from the marine population are similar, suggesting recent common ancestry of the selected alleles in these populations (fig. 3C and table S2). However, several DIs deviate from this pattern to varying degrees. For each DI at each population, we define rigidity as the extent of the overlap between its marker SNPs with those in other freshwater populations (see Materials and Methods for details). We also calculate the average DI rigidity over all populations. The mean rigidity across all DIs is 0.87 (table S2), implying that the majority of marker SNPs coincide between populations (fig. 3D). However, for four of the DIs, rigidity is below 0.4 (figs. 3E-F and table S2). This suggests that for some DIs, the selective sweeps responsible of their origin were soft. The DIs with extremely low rigidity are III-2, which shares only 17% of marker SNPs between the two populations that were used to calculate its rigidity (fig. 3E), and X-2, for which this figure is 6% (fig. 3F). Remarkably, these two DIs overlap genes that are involved in immune response: *CD48* and *SLAM6* in III-2; and *MHC*-associated gene (*mhclzea* (Kersey et al. 2016)) and *CXADR* in X-2. Furthermore, these two DIs possess above average rates of recombination (41.0 and 11.0 respectively, which is higher than the genome average of 8.2).

**Fig. 3.**
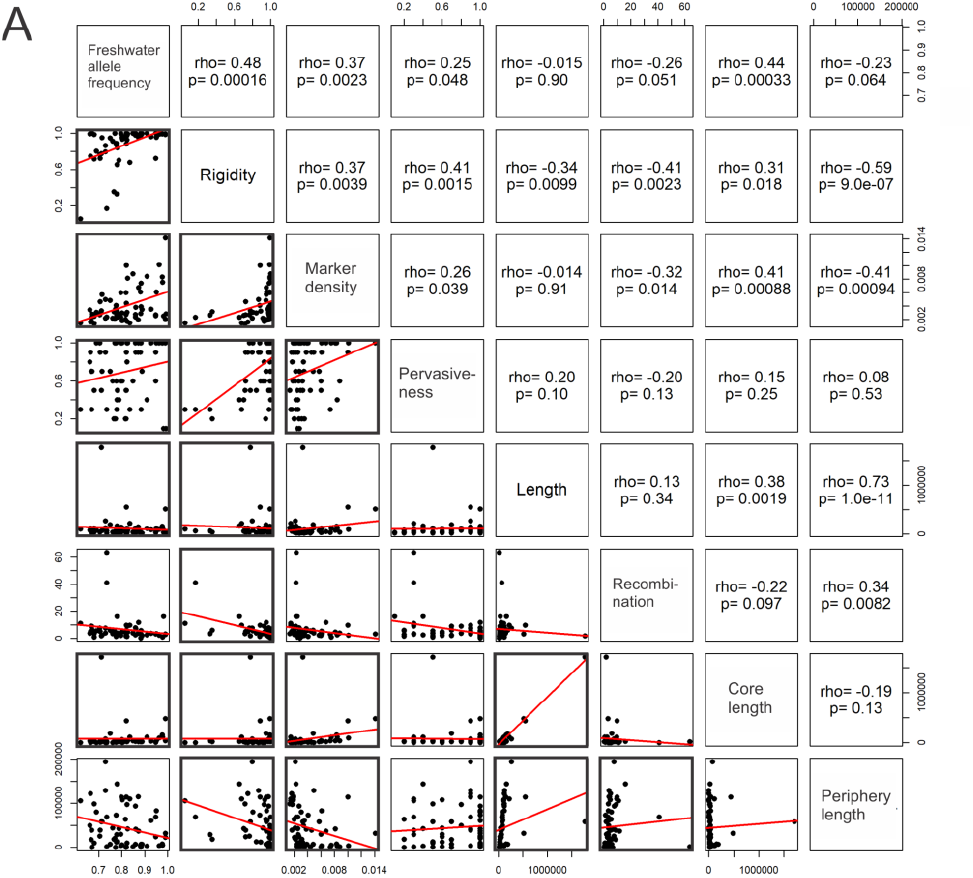
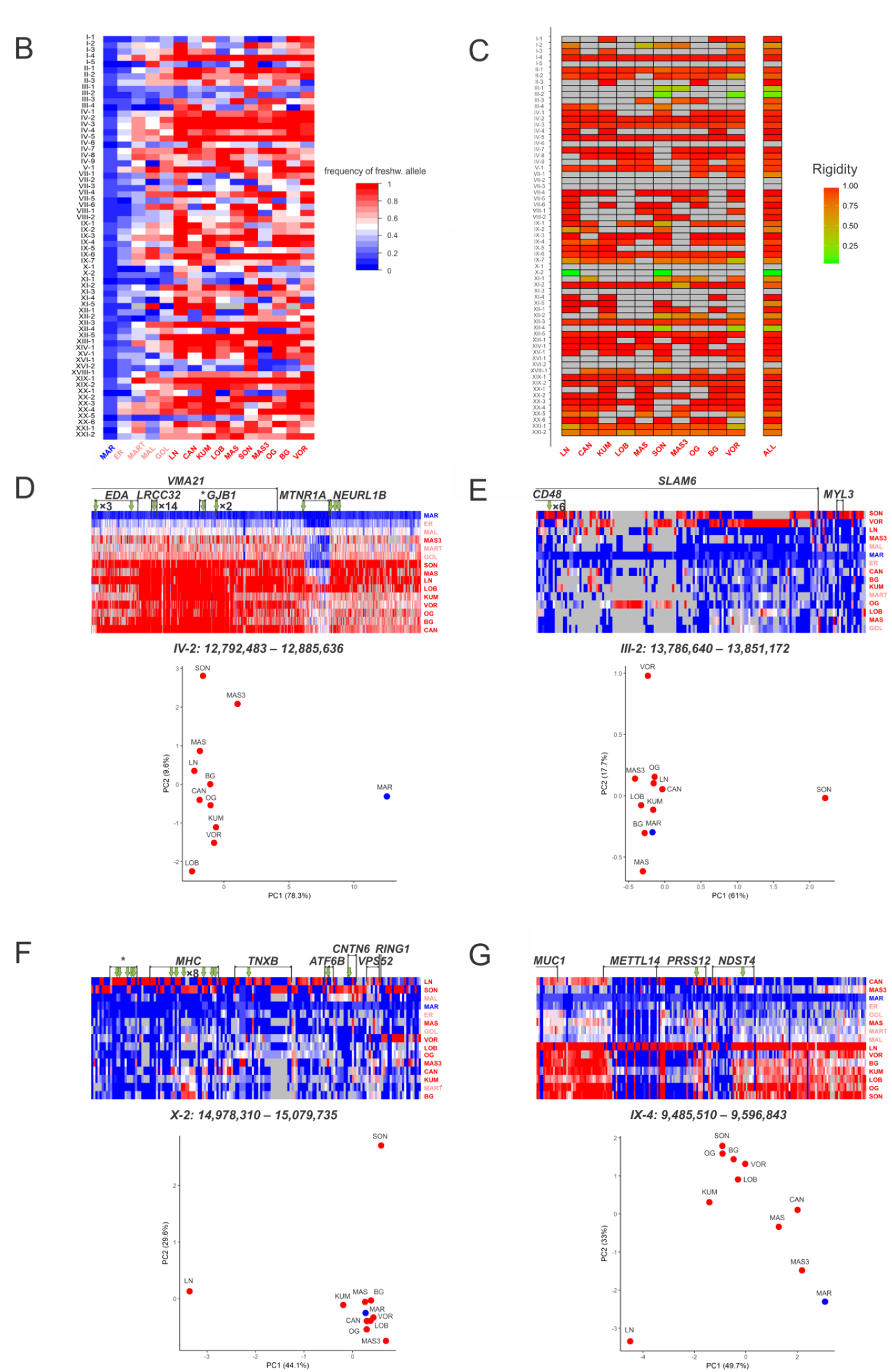
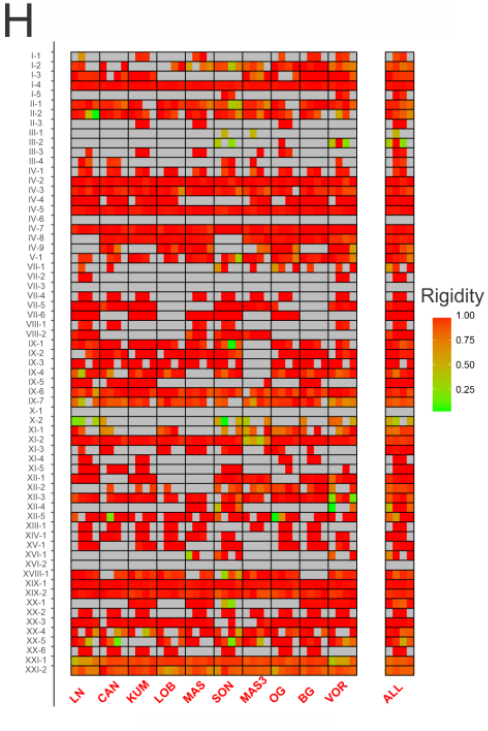
Genomic architecture of the DIs. **(A)** Pairwise scatterplots of different characteristics of the 65 DIs. Black squared frames denote scatterplots with significant Spearman correlations (P < 0.05). **(B)** Freshwater allele frequencies within the 65 identified DIs. Columns, populations from left to right: one marine population, four young freshwater populations and ten freshwater populations of older ages. Color codes for populations: blue – marine; coral – young freshwater; red – older freshwater. Rows, DIs. Grey cells correspond to values that are missing due to insufficient sequencing coverage. **(C)** DI rigidity. Columns, individual populations, or the values averaged over all populations (rightmost column); rows, DIs. Only freshwater populations in which this DI had more than 20 marker SNPs were considered; the remaining cells are colored grey. **(D-G)** Allelic composition of individual DIs. Rows, populations. Columns, marker SNPs within the DI, with genes overlapping the DIs indicated with brackets (unannotated genes are marked with an asterisk); green arrows indicate nonsynonymous marker SNPs. Cell color, freshwater allele frequency. Grey cells correspond to values that are missing due to insufficient sequencing coverage. Scatterplots are PCA plots based on densities of marker SNPs in each population. **(D)** High rigidity (R = 0.99) DI IV-2; most of the marker SNPs are shared between all populations. As a result, the freshwater populations form a single dense cluster along PC 1. Many of the marker SNPs are nonsynonymous, including 14 marker SNPs within a short exon of the *LRRC32* gene. **(E)** Low rigidity (R = 0.17) DI III-2. Most of the marker SNPs with high freshwater allele frequency in the SON population have low freshwater allele frequency in the VOR population, and vice versa. As a result, these two populations are remote in the PCA plot. **(F)** Low rigidity (R = 0.06) DI X-2. **(G)** Variable rigidity DI IX-4; most of the marker SNPs are shared between populations at the 5’ part of the DI, but are private to some of the populations at the central part of the DI. **(H)** The rigidity of a DI varies along its length. This panel is similar to panel B, except that each DI is subdivided into four equal bins along its length, and rigidity is calculated independently for each bin. Only freshwater populations in which this DI had more than 5 marker SNPs were considered; the remaining cells are colored grey.

Multiple locally adaptive haplotypes are unlikely to coexist for a long time unless the selective advantage conferred by them is very similar and the population is highly structured. Therefore, we hypothesized that the DIs with low rigidity tend to be young. In line with this hypothesis, the 6 DIs with the lowest rigidities, which is 10% of all DIs for which rigidity could be calculated, carried fewer marker SNPs per site (0.0022), indicative of their younger age, than the remaining DIs (0.0044; one-sided Wilcoxon test P = 0.01), and the marker density is positively correlated with rigidity across all DIs (ρ = 0.37, P = 0.0039; fig. 3A).

Furthermore, it has been suggested that selection giving rise to soft sweeps should be weaker and/or more local than that giving rise to hard sweeps (Hermisson and Pennings 2005). Therefore, we hypothesized that the DIs with low rigidity will also be less pervasive. Consistently, the pervasiveness of 6 of the DIs with the lowest rigidities is also low (0.3, compared with the 0.8 for the remaining DIs; one-sided Wilcoxon test P = 6.32×10^−5^), and the overall correlation between these two characteristics of a DI was significant (ρ = 0.41, P = 0.0015; fig. 3A).

Finally, soft selection sweeps are more likely to arise under stronger recombination, compared to hard sweeps (Hermisson et al. 2017). Consistently, 6 DIs with the lowest rigidity values had the average recombination rate of 14.5, compared to 4.3 in all other DIs (one-sided Wilcoxon test P = 0.0014), and there was a correlation between recombination and rigidity across all DIs (ρ = –0.41, P = 0.0023; fig. 3A).

After a soft sweep, a particular DI can be represented by a variety of haplotypes. To investigate whether the infrequent haplotypes are different from more frequent ones, we studied the rigidity and marker density of DIs in each population. As expected, the 10% of haplotypes with the lowest rigidity values, i.e. those that deviate most from other haplotypes in the same DI (see Materials and Methods), tend to be younger than the remaining haplotypes (mean densities of marker SNPs: 0.0019 vs. 0.0041, one-sided Wilcoxon test P = 2.58×10^−9^; fig. S1A), and also have larger periphery regions (48.0 Kb vs. 29.2 Kb for DIs excluding inversions, one-sided Wilcoxon test P = 0.0025; fig. S1B).

### Evolutionary history of a DI may vary along its length

We hypothesized that different segments of an individual DI may differ in their evolutionary history. To study this, we have first subdivided each DI into four bins along its length, and analyzed these bins independently. For some of the DIs, we found that rigidity varies along their length (figs. 3G–H). This implies that even within a single DI, some segments may be involved in hard selective sweeps, while other segments are involved in soft sweeps. Similar to the DI-level analysis, the low-rigidity segments of DIs usually also have low densities of marker SNPs, indicating that they are younger. This is true both for the rigidity averaged across all populations (0.00230 for the 18 segments with the lowest rigidity, which is 10% of all considered segments, compared to 0.00495 for the remaining segments; one-sided Wilcoxon test, P = 0.0021; fig. S2A) and when rigidity is calculated for each population individually (0.00068 vs. 0.00380 for the 10% of all considered segments versus all the remaining segments; one-sided Wilcoxon test P < 2.2×10^−16^). The segments with low average rigidity were also less pervasive (0.53 vs. 0.72 respectively, one-sided Wilcoxon test P = 0.0036; fig. S2B). Therefore, some of the DIs were likely comprised of an older segment resulting from an ancient hard sweep, neighbored by a younger segment resulting from a soft sweep (fig. 3H). Finally, the low-rigidity segments were characterized by a higher recombination rates than the remaining segments (9.94 vs. 5.46, one-sided Wilcoxon test P = 6.18×10^−4^; fig. S2C). Therefore, the differences between parts of a DI likely result from the differences in haplotype ages and from the underlying recombination structure.

The 25% length bins of the same DI sometimes also had radically different values of pervasiveness: while the markers contained near the center of the DI carried freshwater alleles in all or nearly all freshwater populations, near the DI edges, some of the populations were comprised solely of marine alleles (e.g. fig. S3).

To study this in more detail, we defined population-level DIs for each population independently, and studied the reproducibility of their boundaries between populations. In general, the coordinates of the population-level DIs matched well between populations, or at least overlapped strongly. The positions of their boundaries were similar: the boundaries of 65% of population-level DIs where within 50 Kb of the boundaries of the DI defined from all populations; 84%, within 100 Kb; and the rest 16% on the distance within 200 Kb (table S5)). For the three inversion DIs, the positions of the boundaries overlapped by 79% across populations as expected. However, even in some of the DIs not associated with inversions, e.g., IV-3, V-1, the boundaries coincided precisely between some of the populations in which they fired (figs. S3A–B). Despite the overall high conservation of the positions of the boundaries, in some population pairs, the population-level DIs comprising the same DI overlap only marginally; DIs II-2, XVIII-1 are examples of this (figs. S3C–D).

For each DI, let us call the segment of the genome which is included in all population-level DIs its core part, and the remainder of the DI, its periphery. The length of the periphery is not correlated with the length of the core (ρ = −0.19, P = 0.13; fig. 3A). By contrast, it is negatively correlated with the DI rigidity (ρ = –0.59, P = 9.0×10^−7^), which implies that similar forces could contribute to the diversity of allelic composition and the lengths of haplotypes. The recombination rate is higher on the periphery compared with the core region (average recombination rate for core and peripheral segments are 5.39 and 7.15 respectively, one-sided Wilcoxon test P =0.008). This could be in part due to variation in the rate of recombination, and the recombination rate is correlated with rigidity (ρ = −0.41, P = 2.3×10^−3^) and periphery length (ρ = 0.34, P = 8.2×10^−3^; fig. 3A).

The density of marker SNPs, which is indicative of the DI age, is positively correlated with the DI core length (ρ = 0.41, P = 8.8×10^−4^ (fig. 3A)). This is consistent with the divergence hitchhiking model, which predicts that a DI should expand with time as novel marker SNPs arise at genomic regions adjacent to it.

### Putative balancing selection on DIs

Because we focused on marker SNPs with very different frequencies between the marine and the freshwater populations, in the majority of the detected DIs, the frequency of the freshwater haplotype in the freshwater populations is high. However, this is not always the case. A striking exception is the DI XXI-1 which coincides with the longest identified inversion (Jones et al. 2012). In this DI, in the marine population, the frequency of freshwater alleles in marker SNPs is very low (~4%), which is lower than for an average DI (~11%). In all freshwater populations, the frequencies of freshwater alleles are elevated; however, contrary to what we see in most other DIs, they always remain at an intermediate level and never reach 100% (the mean allele frequency across all freshwater populations: 0.54, range: 0.29-0.81; table S1). This could be explained by a weaker positive selection on this DI in freshwater populations. However, under moderate selection, we would expect a strong dependence of the freshwater haplotype frequency on the population age (Terekhanova et al. 2014). For DI XXI-1, we see no such dependence. Moreover, in the two very young freshwater populations from our previous study (Terekhanova et al. 2014), the freshwater haplotype frequency in this DI is already rather high (43% and 65% after 30 and 250 years, respectively).

Intermediate allele frequencies in DI XXI-1 in multiple freshwater populations are also consistent with balancing selection. One form of balancing selection is heterozygote advantage. To test whether heterozygotes are overrepresented at this locus, we performed allele-specific PCR for this locus in 9 individuals from the Mashinnoye lake (MAS), and in 7 individuals from the Lobaneshskoye lake (LOB) from the previous study (Terekhanova et al. 2014). The deviation from the Hardy-Weinberg proportions was observed for one population (chi-square test, MAS: P = 0.16, LOB: P = 3.5×10^−9^, table S6).

In some other DIs, the freshwater allele frequency is also intermediate and independent of the age of the population (table S1). As candidate targets of balancing selection, we have selected the DIs with the lowest difference in freshwater allele frequencies between the young and old freshwater populations. The first four DIs on the list were IV-8, XXI-1, IX-7, IV-4. The first three of these, IV-8, XXI-1 and IX-7, overlap multiple genes involved in the immune system: DI IV-8 overlapped with *HSPA9* and positioned within 15 Kb of the *IGBP1* and *MAGT1;* the large inversion DI XXI-1 overlapped with *RBCK1, SOCS6, CD226, RRS1;* and DI IX-7 overlapped with *CLEC6A, CD209* and *UNC93B1*. The fourth DI IV-4 overlapped a pair of duplicated *AKR1B1* genes involved in reproduction (table S4).

### Positive selection outside DIs

Previously, we have shown that marker SNPs are enriched in nonsynonymous substitutions compared with the non-marker SNPs segregating within the marine population (Terekhanova et al. 2014), and interpreted this as a sign of positive selection. Here we analyze marker SNPs using data on many populations. In line with our previous results (Terekhanova et al. 2014), we find that the dN/dS ratio for marker SNPs outside DIs is higher than that for marker SNPs within DIs, signifying high prevalence of positive selection outside DIs (fig. 4). Notably, the dN/dS ratio for marker SNPs is higher for those marker SNPs that are present in at least five populations, compared with all other marker SNPs, both inside the DIs (0.301 vs. 0.243) and outside them (0.615 vs. 0.315; fig. 4). Therefore, the fraction of positively selected marker SNPs is the highest among those marker SNPs at which the freshwater allele is present in many populations.

**Fig. 4.**
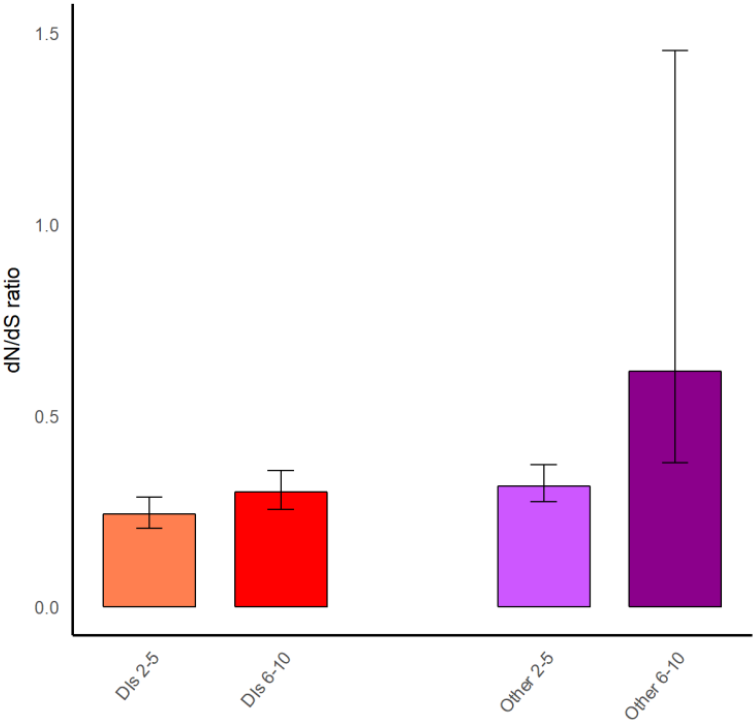
dN/dS ratio for marker SNPs. dN/dS ratios calculated for marker SNPs which are found in 2-5 freshwater populations (left bar in each pair) or in the 6-10 freshwater populations (right bar in each pair). Red bars correspond to the ratios calculated for marker SNPs located in DIs, and purple bars, to the ratios calculated for marker SNPs located outside of the DIs.

## Discussion

The high speed of adaptation of threespine stickleback populations to the freshwater environment is made possible by the fact that the freshwater alleles are present at low frequencies in the ancestral marine population (Colosimo et al. 2005; Schluter and Conte 2009). Adaptation to such a radically different environment is likely to be genetically complex and to involve many loci, as was shown for other species (Renaut et al. 2013; Gao et al. 2017). Identifying all loci responsible for a complex adaptation is a difficult task (Hoban et al. 2016). However, in threespine sticklebacks, similar to other species that have adapted to widely different environments (Jones et al. 2012; Nadeau et al. 2012; Renaut et al. 2013; Sodeland et al. 2016), some of the loci responsible for adaptation are located in DIs – regions of elevated divergence between the freshwater and marine populations (Turner et al. 2005; Feder and Nosil 2010). We do not know what proportion of adaptive differences between the marine and freshwater populations of threespine stickleback are confined to DIs, although this proportion is likely to be high (Terekhanova et al. 2014).

DIs are scattered through the genome, and are relatively easy to identify as sufficiently long regions with an increased density of marker SNPs – sites where marine and freshwater populations carry different common alleles. It is not clear what factors promote DIs formation and are responsible for variation in their lengths. A relatively long DI may arise due to multiple targets of positive selection located within a relatively short genomic region, to very strong selection acting on just a single target (Feder et al. 2012; Flaxman et al. 2013), or to locally reduced recombination rate (Feulner et al. 2015; Samuk et al. 2017).

To elucidate the processes involved in DIs formation, we studied ten independent freshwater populations of threespine stickleback which originated recently in the basin of the White Sea. We found that DIs tend to reside in genomic regions of low recombination rate, in line with the previous observations (Samuk et al. 2017), probably because reduced recombination facilitates their formation (Barton 2000; Yeaman et al. 2016). This may seem paradoxical because recombination usually facilitates adaptation (Felsenstein 1974).

However, low recombination rate also makes adaptation easier to detect by increasing the length of a DI which emerges as a result of positive selection acting on an individual target (Jones et al. 2012; Nadeau et al. 2012; Renaut et al. 2013; Sodeland et al. 2016). Reduced recombination is not a necessary condition for the formation of DIs: around some of them, recombination is very fast (table S2).

DIs also tend to be clustered along chromosomes, and this effect cannot be explained by differences in the recombination rate. The clustering of DIs was previously observed in the Atlantic cod (Bradbury et al. 2013) and cichlid species (Malinsky et al. 2015); DIs also seem to be clustered in stick-insects (Riesch et al. 2017) and munias (Stryjewski and Sorenson 2017), although no statistical analysis has been performed. In human populations, some of the genomic regions that likely harbored selective sweeps, as defined by the iHS scan, are also clustered along the chromosomes, and this clustering can be only partially explained by variation in recombination rate, gene density, or background selection (Johnson and Voight 2018).

Possibly, DIs may be clustered because neighboring DIs facilitate formation of each other, for example, in the process of divergence hitchhiking. This process increases the probability of fixation of a new beneficial mutation located near another beneficial mutation (Via 2009; Feder et al. 2012), thus expanding a DI or producing distinct tightly linked DIs. As a result, when two incompletely isolated populations adapt to different environments, the locally adaptive alleles tend to reside in tightly linked loci, forming long haplotype blocks (Yeaman and Whitlock 2011; Yeaman et al. 2016). Consistently with the divergence hitchhiking model, we find that older DIs which possess higher densities of marker SNPs tend to have longer core region shared between populations (fig. 3A). A similar pattern has been observed in cichlid species pairs, where more differentiated species have more diverged DIs (McGee MD et al. 2015), and in lake whitefish species pairs, where the most differentiated pairs of species have longer DIs (Renaut et al. 2012).

Under divergence hitchhiking, one may also expect to see similar frequencies and positive LD between freshwater alleles at adjacent DIs. However, this prediction of the model is not confirmed by our data: frequencies of freshwater alleles in nearby DIs are no more similar than in remote DIs (fig. 3B and table S1). Similarly, in previous studies, positive LD was observed only for a few of the adjacent DIs (Hohenlohe et al. 2011); and DI divergence and length were found to be independent of the age of the locally adapted population (in the range of thousands of years) (Feulner et al. 2015). This discrepancy is perhaps not surprising. While the attraction of the DIs may be manifested at timescales of DIs lifespan which may cover millions of years (Nelson and Cresko 2018), its signal may be too weak to be detected at timescales of individual populations which are only thousands of years old.

The number of DIs responsible for the adaptation of threespine stickleback to freshwater that have been detected throughout its range is in the high tens (Hohenlohe et al. 2010; Jones et al. 2012; Terekhanova et al. 2014). Although the set of DIs is far from being identical across populations, often some of these DIs are reused by freshwater populations of independent origin. It seems plausible that some of the DIs are particularly important for adaptation, and they can be expected to be older and more pervasive. Indeed, pervasive DIs possess greater density of marker SNPs, carry freshwater alleles at higher frequencies in freshwater populations, and have longer core region shared between populations (fig. 3A and table S2). Interestingly, the frequency of the freshwater allele in pervasive DIs tends to be higher than in other DIs even in the marine populations (fig. 3B), suggesting that the selection against the freshwater alleles in the marine environment at such DIs can be weaker. The elevated frequencies of freshwater alleles in the pervasive DIs in the ancestral marine population can facilitate their frequent fixation in freshwater populations. Indeed, the frequencies of freshwater alleles in pervasive DIs are higher than in other DIs even in the youngest freshwater populations (correlations between pervasiveness and freshwater allele frequency: lake Ershovskoye (ER), 30 years old (Terekhanova et al. 2014), ρ = 0.58, P = 4.33×10^−7^; lake Martzi (MART), 250 years old (Terekhanova et al. 2014), ρ = 0.63, P = 1.42×10^−8^; fig. 3B).

The older a DI, as revealed by its marker density, the more rigid it generally is. This can mean that old DIs originated through hard sweeps, while younger DIs sometimes originated through soft sweeps. Alternatively, the correlation between the DI age and rigidity can arise even if the ratio of hard to soft sweeps at their origin was the same, because over the long lifetime of a DI, multiple haplotypes resulting from a soft sweep are more likely to be outperformed or lost due to random drift. This latter scenario is supported by some theory (Yu and Etheridge 2010) and data (Lee and Marx 2013; Anderson et al. 2017). Most DIs we detect are relatively old and rigid, with average rigidity R = 0.87. Note that we would not be able to detect a very young DI, since its marker density would not pass our thresholds for detection.

Although the average rigidity of a DI is high, it is below 0.4 for four of them. This suggests that multiple haplotypes were involved in adaptation at a single DI (Bassham et al. 2018), i.e., that sweeps at their origin were soft. Two kinds of soft sweeps are possible. Exactly the same beneficial allele can arise against multiple backgrounds. Alternatively, a soft sweep can involve selection of different, although functionally similar, beneficial alleles (Hermisson and Pennings 2005). Because we are unable to precisely identify the targets of positive selection in our DIs, we cannot say, for a DI of low rigidity, which kind of soft sweep has led to its origin. Still, non-rigid DIs usually harbor some proportion of common marker SNPs: even the least rigid DI shares 6% of the marker SNPs between the only two populations in which it has fired (DI X-2, fig. 3F). Therefore, we cannot reject the simplest hypothesis that the beneficial allele involved in adaptation in a DI has been exactly the same in all populations. Some of the least rigid DIs are characterized by above average recombination rates, implying that they also have elevated local effective population sizes (*Ne*) (Gossmann et al. 2011), probably because they possess genes in which diversity and recombination are beneficial, such as immune and signaling pathways genes (The International HapMap Consortium 2007; Choi et al. 2016). Also, the presence of multiple haplotypes, with traces of recombination between them, in regions of increased divergence among multiple populations could also result from differential sorting of ancestral variation as has recently been observed in munia species (Stryjewski and Sorenson 2017). We find that the architecture of a DI may vary along its length (figs. 3H and S1), implying that different types of sweeps could have played a role in formation of even a single DI. At this within-DI level, segments with the lowest rigidities tend to be younger and possess lower densities of marker SNPs, similar to what is observed for individual DIs.

According to our criterion for identification of DIs (see Materials and Methods), coordinates of a DI in individual populations do not need to overlap. However, we find that usually these coordinates overlap substantially, so that a DI possesses a long shared core region. We also see that the length of the periphery of a DI sometimes varies only slightly. Since most DIs are old, this implies that recombination within a DI may be constrained. Such a constraint could arise due to strong divergent selection and/or to structural variation. The high conservation of DI boundaries over millions of years of their evolution is in line with the theoretical prediction that DIs should accumulate genomic re-arrangements that maintain their lengths (Yeaman 2013). Indeed, three of the analyzed DIs reside within inversions which impede recombination (Jones et al. 2012). Linkage is also increased inside the non-inversion DIs ((Hohenlohe et al. 2011), fig. 1C), which promotes stability of their boundaries, and even between some of them (Hohenlohe et al. 2011). Still, some DIs have only short core regions or even do not overlap at all (fig. S3).

In general, selection acting at a DI is strong: the average frequency of freshwater alleles across the fired DIs is 0.81 (table S2). However, in some cases, we observe that the freshwater alleles at a DI only reach an intermediate frequency. This could be due to two reasons: the selection in favor of the freshwater alleles at this DI is weak, so that they have had insufficient time to reach a high frequency; or the equilibrium allele frequency is below 1 due to the action of balancing selection. The first explanation for our observations is unlikely, because the selection coefficients for favorable alleles at DIs are usually very high and because the freshwater allele frequency at these DIs is independent of the age of the population (Barrett et al. 2008; Terekhanova et al. 2014). By exclusion, we are left with the second scenario, although it is difficult to test with our data. Individual genes, especially those involved in immune response, may experience balancing selection even within a single habitat due to mechanisms such as heterozygote advantage, frequency-dependent selection, or fluctuating selection. The action of balancing selection in the evolution of immune genes is supposed to be the result of host-parasite interrelations (Eizaguirre et al. 2012).

The top candidate for balancing selection is DI XXI-1. It is contained within the longest inversion and carries an unusually high density of marker SNPs (Terekhanova et al. 2014), suggesting its old age at the level of the metapopulation, and it was inferred to be one of the oldest among all DIs (~8 Mya (Nelson and Cresko 2018)). Although intermediate frequency of the freshwater haplotype and independence of this frequency on age suggests that this DI could have experienced balancing selection, we observe the deviation from the Hardy-Weinberg equilibrium only in one of two populations tested. Instead, the selection involved could be negative frequency-dependent, as reported recently for the major histocompatibility complex class IIβ (MHC IIβ) genes in stream-lake stickleback populations (Bolnick and Stutz 2017).

## Materials and Methods

### Collection of samples and ethical statement

We analyzed 10 independent freshwater populations of older age, the youngest of which is ~600 years old, together with 4 younger populations, the oldest of which is ~250 years old (table 1). The 4 younger populations and two of the older populations (MAS and LOB) have been analyzed previously (Terekhanova et al. 2014). Two interconnected Kumyazh’i lakes were pooled into one sample (KUM). We also obtained marine individuals from a novel location (White Sea, WSBS) and pooled them with the two previously analyzed marine samples (Nilma and anadromous individuals from lake Ershovskoye). We collected 8-24 fish from each population, which were then pool-sequenced at the average coverage of 36x (table 1).

Fish were caught by scoop-net or landing-net, anaesthetized and euthanized with a tricaine methane sulphonate solution (MS222), and then immediately fixed in 96% ethanol. Fish collection was conducted under the supervision of the Ethics Committee for Animal Research of the Koltzov Institute of Developmental Biology Russian Academy of Sciences.

### Genome sequencing

Total genomic DNA was extracted from each individual using Wizard genomic DNA purification kit (Promega). Prior to library preparation, DNA samples of between 8 and 24 (table 1) fish from the same population were pooled in equal proportions. Samples from populations OG, BG, VOR, MAS3 and SON were prepared with TruSeq PCR-free protocol (Illumina) and sequenced using HiSeq4000 with 150 bp paired-end reads at Norwegian Sequencing Center (Oslo, Norway). The remaining samples WSBS, KUM, LN and CAN were prepared according to the TruSeq DNA Sample Preparation Guide (Illumina) and sequenced using HiSeq2000 with 101 bp paired-end reads. Sequencing reads for each population are available at the NCBI Short Read Archive (http://www.ncbi.nlm.nih.gov/sra; accession numbers of the projects are SRP023197 and SRP151980).

### Genome mapping

The reads were trimmed with trimmomatic version 0.27 (Bolger et al. 2014) and then mapped to the reference genome of the *G. aculeatus* obtained from the UCSC using bwa mem program from the BWA package (Li and Durbin 2009). The alignment was then converted to bam and sorted with the programs from the samtools package (Li 2011). Aligned reads were processed with picard-tools (http://broadinstitute.github.io/picard/) to remove duplicate reads. SNPs were called with the mpileup program of the samtools package (Li 2011).

### Identification of DIs

To identify the DIs, we used the 10 freshwater populations of older age (table 1). For each population, we first traversed the genome in 10 Kb genomic sliding windows (in 1Kb steps), listing all windows carrying at least 10 marker SNPs, and calculated the average frequency of the freshwater allele at marker SNPs in those windows. Next, from those windows, we picked the window with the maximum mean freshwater allele frequency; if the freshwater allele frequency in this window was > 0.5, this window was then considered the “seed” of a putative population-level DI. We then extended this putative DI along the sequence in each direction by merging it with adjacent 10 Kb windows in 1 Kb steps, until a window was reached in which the mean freshwater allele frequency has declined by more than 30%, compared to the seed window. Then the window with the next highest freshwater allele frequency was used, and the procedure was repeated till no more seeds could be identified outside the already identified putative population-level DIs. After repeating this procedure for all populations, we merged all putative population-level DIs across all populations with lengths of 15 Kb or more if they were located within 30 Kb from each other to obtain the putative list of DIs. For the final list of DIs, we kept only those putative DIs containing at least one 10 Kb window with more than 50 SNPs, each of which was a marker SNP in at least one population. The freshwater allele frequency of a DI at a given population was defined as its mean frequency across the 20% of the windows with the highest freshwater allele frequency within the DI. Finally, to obtain the ultimate list of population-level DIs, we merged all putative population-level DIs within the boundaries of a single DI, irrespective of the distance between them; and discarded those population-level DIs that had not fired, i.e., where the freshwater allele frequency was below 0.5.

### Pairwise correlations between populations

Pairwise correlation analysis was performed on the 10-coordinate vectors of individual DIs, where each coordinate corresponds to the freshwater allele frequency in each of the ten older freshwater populations studied. P values were obtained with the usage of the MPT.Corr.R (Yoder et al. 2004). PCA-analysis was performed with R. Freshwater allele frequency was calculated as the mean across the 10 Kb windows with the highest values covered 20% of the DI.

### Fst, Dxy and π calculation

We calculated Fst, Dxy and π values for all 5 Kb non-overlapping genomic regions. To calculate Fst, we used mpileup2sync.jar and fst-sliding.pl programs from the popoolation2 package (v. 1201) (Kofler et al. 2011). We calculated Dxy and π as the average value across all sites with 1 or 2 alleles. Dxy was calculated at each site as p_11_ × p_22_ + p_12_ × p_21_, where p_11_ and p_12_ are the frequencies of the two alleles in freshwater population and p_21_ and p_22_ are the frequencies of the same alleles in marine population. π at each site was calculated as 1 – (p_1_^2^ + p_2_^2^), where p_1_ and p_2_ are the frequencies of the two alleles.

### Genomic annotation and GO analysis

Genomic annotation was obtained from the Ensembl database release 72 (Ruffier et al. 2017). We also used BLAST search (blastx algorithm) against the nr database to annotate stickleback genes overlapping DIs, using the top BLAST result hit as the homolog if its e-value was below 0.01. GO analysis was performed using the R package clusterProfiler (Yu et al. 2012).

### Recombination rate

We obtained the mean coalescent-based population recombination rates for each 10 Kb window in the G2L freshwater lake population from (Feulner et al. 2015) (Philine G. D. Feulner, personal communication). These recombination rates are population size-scaled, ρ = 4*N_e_r*, where *r* is the number of expected cross-over events per 10 Kb per generation. For the permutation analyses, the mean values over all 10 Kb windows were obtained for each 1 Mb window, and the values were categorized into five bins. Each DI was classified as belonging to one of the categories; if it fell on a boundary between two bins, we recategorized the genomic segment carrying the smaller part of the DI as belonging to the same bin as the larger part of the DI, so that each DI would fall into one bin. Permutation analysis was performed with shuffleBed from the bedtools2 (Quinlan and Hall 2010) with the additional parameter –noOverlapping. P values were obtained from 1000 independent permutation trials.

### Rigidity

We calculated rigidity for each DI as follows. We chose those populations possessing at least 20 marker SNPs. For each of the *N* pairs of such populations, we determined the corresponding sets of marker SNPs, *A* and *B*, i.e. SNPs with the frequency of the freshwater allele > 0.8 (see above) in the corresponding population. To avoid misclassifying a shared marker SNP as present in just one of the two compared populations, if it was present in one of the populations with frequency ≥ 0.8, but in the other, with the frequency between 0.5 and 0.8, it was also considered present in both populations; this correction may bias rigidity estimates upwards. We then calculated rigidity as the mean, over all *N* pairs, of the ratio of the number of the SNPs shared between *A* and *B* and the lesser of the numbers of SNPs in *A* and *B*:

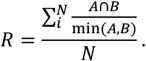

The rigidity for the 25% genomic segments of DIs were calculated similarly, except that we required presence of 5, rather than 20, marker SNPs in a segment.

The rigidity of a DI for a particular population was estimated similarly, except that only those population pairs involving the considered population were used.

## Acknowledgements

We thank Maria D. Logacheva and Aleksey A. Penin for help with library preparation; Philine G. D. Feulner for providing the coalescent-based population recombination rates; and Lubov Mugue and Andrey Bulakhov for participating in sample collection.

